# How Axon and Dendrite Branching Are Governed by Time, Energy, and Spatial Constraints

**DOI:** 10.1101/2021.07.15.452445

**Authors:** Paheli Desai-Chowdhry, Alexander Brummer, Van Savage

## Abstract

Neurons are connected by complex branching processes - axons and dendrites - that collectively process information for organisms to respond to their environment. Classifying neurons according to differences in structure or function is a fundamental part of neuroscience. Here, by constructing new biophysical theory and testing against our empirical measures of branching structure, we establish a correspondence between neuron structure and function as mediated by principles such as time or power minimization for information processing as well as spatial constraints for forming connections. Specifically, based on these principles, we use undetermined Lagrange multipliers to predict scaling ratios for axon and dendrite sizes across branching levels. We test our predictions for radius and length scale factors against those extracted from neuronal images, measured for cell types and species that range from insects to whales. Notably, our findings reveal that the branching of axons and peripheral nervous system neurons is mainly determined by time minimization, while dendritic branching is mainly determined by power minimization. Further comparison of different dendritic cell types reveals that Purkinje cell dendrite branching is constrained by material costs while motoneuron dendrite branching is constrained by conduction time delay over a range of species. Our model also predicts a quarter-power scaling relationship between conduction time delay and species body size, which is supported by experimental data and may help explain the emergence of hemispheric specialization in larger animals as a means to offset longer time delays.

**Author summary:** Neurons are the basic building blocks of the nervous system, responsible for information processing and communication in animals. They consist of a centralized cell body and two types of processes - axons and dendrites - that connect to one another. Previous studies of the differences among neuron cell types have focused on comparisons of either structure or function separately, without considering combined effects. Based on theory for structure of and flow through biological resource distribution networks, we develop a new model that relates neuron structure to function. We find that differences in structure between axons and dendrites as well as between dendrites of different cell types can be related to differences in function and associated evolutionary pressures. Moreover, using our mathematical model, we find that the conduction time delay of electrical signals systematically varies with species body size - neurons in larger species have longer delays - providing a possible explanation for hemispheric specialization in larger animals.

## Introduction

Neurons are fundamental structural units of information processing and communication in animals. They are made up of a centralized cell body, called the soma, and two types of extending processes, axons and dendrites. These processes transfer information between cells in the form of electrical and chemical signals. Axons generally conduct signals from the cell body to the synapses, where they connect with the dendrites of other neurons. These dendrites generally conduct signals from the synapse to the cell body. The processes form synaptic connections with one another in complex patterns. Different types of cells exhibit diverse morphological forms - some neurons have no axons or dendrites, while some have long axon processes that extend over meters, and others have vast dendritic trees that branch extensively to fill two- or three-dimensional space, corresponding to the mathematical and modeling concept known as space-filling [Johnston and Wu, 1995].

Seminal studies in neuroscience characterized morphological differences across cell types. For instance, Santiago Ramón y Cajal’s “Histology of the Nervous System of Man and Vertebrates” is considered to be the founding document of neurobiology [Zeng and Sanes, 2017], consisting of detailed drawings and comparative descriptive analysis of neuron morphology across different cell types and species [Ramón y Cajal, 1995]. Modern techniques and devices have allowed for more precise quantitative measurements at the single-cell level. Indeed, recent work has established quantitative morphological distinctions across different cell types, focusing on quantities such as mean dendritic length, total dendritic length, and number of branching points [Gertler et al., 2008 and Lu and Yang, 2017].

As vast as the structural diversity is, there is an even greater diversity of functional properties [Johnston and Wu, 1995]. Within sensory, motor, and interneurons, there are different types of neurotransmitters and receptors that affect the nature of signal processing [Squire et al, 2013]. A major future goal of neuron cell-type classification is to establish a correspondence between morphological and functional properties [Zeng and Sanes, 2017]. Here, we seek to address the question of how structural properties relate to neuron function, and whether there are evolutionary driving forces that dictate how morphology is optimized by biological principles or pressures.

A promising approach to the relationship between neuron structure and function is biological scaling theory, as it has previously been applied to understand patterns in the branching structures of biological resource distribution networks. Generally, a biological property Y scales with body mass M as *Y* = *Y*_0_*M^b^*, where *Y*_0_ is a proportionality constant and b is a scaling exponent [West et al., 1997]. An example is metabolic rate, scaling with body mass to the power 3/4, a result known as Kleiber’s Law [Kleiber, 1932].

West, Brown and Enquist (WBE) proposed that Kleiber’s law and other biological scaling laws arise because biological organisms are sustained by resource distribution branching networks that are optimized to supply all parts of the body [West et al., 1997]. Past work on cardiovascular networks has employed WBE theory to derive scaling laws for the vessel radius and length as a result of minimizing power loss for fluid flow along with space filling in order to fuel whole organism metabolism [Savage et al., 2008]. Moreover, previous results have shown a quarter-power allometric scaling relationship between cell size and body size in a range of cell types in mammals, including neurons [Savage et al., 2017].

Single neuron cells have structural similarities to cardiovascular networks, with centralized cell bodies analogous to the heart and branching processes analogous to blood vessels. We propose that the branching structures of axons and dendrites result from optimizing organismal function subject to biophysical constraints. We consider biophysical properties of neurons that might play an important role in governing structure, using data to guide our evaluation of the relative importance of different functions.

An important evolutionary function of neuronal networks involves transferring large amounts of information between brain regions in a short amount of time [Laughlin and Sejnowski, 2003]. On the individual cell level, the various morphological forms observed in neurons are various adaptations of basic principles such as limiting signal time delay [Ramón y Cajal, 1995]. Thus, it is important to consider conduction time as an important design principle that governs neuronal branching structures.

Indeed, foundational work by Cuntz et al. has used graph theory to quantify and study how connections among axons and dendrites determine conduction time delay. This approach focuses on the tradeoff between wiring costs and conduction time, represented as path length [Cuntz et al., 2010]. The results formalize the laws set forth by Ramon y Cajal, leading to a graph-theoretical algorithm that generates biologically accurate synthetic axonal and dendritic trees [Cuntz et al., 2011].

However, a key aspect absent from this formalism is that it does not include the diameter of axonal and dendritic fibers nor does it incorporate the principle of space-filling. Axon and dendrite radius relate to resistance and thus signaling speed and conduction time. Space-filling constrains the possible connections, branching, and network structure of neurons. Consequently, in this paper we take a similar approach to Cuntz et al. [Cuntz et al., 2010, Chklovskii, 2004] but now incorporate the dependence of conduction time on fiber radius and myelination - insulation that surrounds the fiber and facilitates signal transduction [Squire et al., 2013] - along with constraints of space-filling.

As the speed of information processing increases, energy loss due to dissipation also increases [Laughlin and Sejnowski, 2003]. Signaling in the brain consumes a substantial amount of energy [Attwell and Laughlin, 2001], which suggests that energy expenditure is another important factor constraining the design. Moreover, the relationship between metabolic rate and conduction time plays an important role in determining axon function in species across scales of body size [Wang et al., 2008] The WBE framework relies on the assumption that resource distribution networks are optimized such that the energy used to transport resources is minimized [West et al., 1997], specifically, by minimizing power lost to dissipation in small vessels [Savage et al., 2008].

Building on biological and physical principles that constrain electrophysiological signaling and information processing in neurons, we build models that predict a suite of neuron morphologies based on which biological or physical principle is under the strongest selection or pressure. Our model includes both conduction time and energy efficiency while also incorporating additional factors set forth by Ramón y Cajal’s laws such as the material costs and space filling [Ramón y Cajal, 1995]. We make theoretical predictions for how branch radius and length change across branching generation for both axons and dendrites. We compare these predictions to our empirically measured data to make conclusions about the functional basis for morphological differences observed across cell types. We also use this model to predict how conduction time delay in neurons changes with neuron size, another prediction that is supported by empirical data.

## Theory

### Model

Because conduction time delay and power usage are fundamental and costly for information processing in neurons, we develop a mathematical objective function to minimize them [Boyd and Vandenberghe, 2004]. Evolutionary pressures and developmental processes shape branching networks and materials (such as myelination) to achieve this.

Equation 1 is a general form of our objective function. The biophysical constraints are represented as arbitrary functions and added to the expressions to be minimized, allowing us to use the method of undetermined Lagrange multipliers to optimize this function.

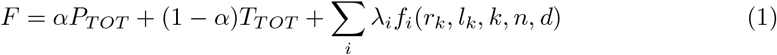

Here, *P_TOT_* is the power lost due to dissipation, *T_TOT_* is the time delay for a signal travelling across the network, r is the branch radius, l is the branch length, k is the branching generation of the network (with 0 being the trunk and N being the tips), N is the total number of levels of the network, n is the branching ratio, M is the mass of the neuron process, and d is the dimension of space into which the neuron processes project. The branching ratio, n, is equal to 2 for a bifurcating network. We use optimization methods to calculate scaling relationships between the radius and length of successive branches, 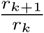 and 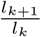, as shown in Figure 1.

**Fig 1.**
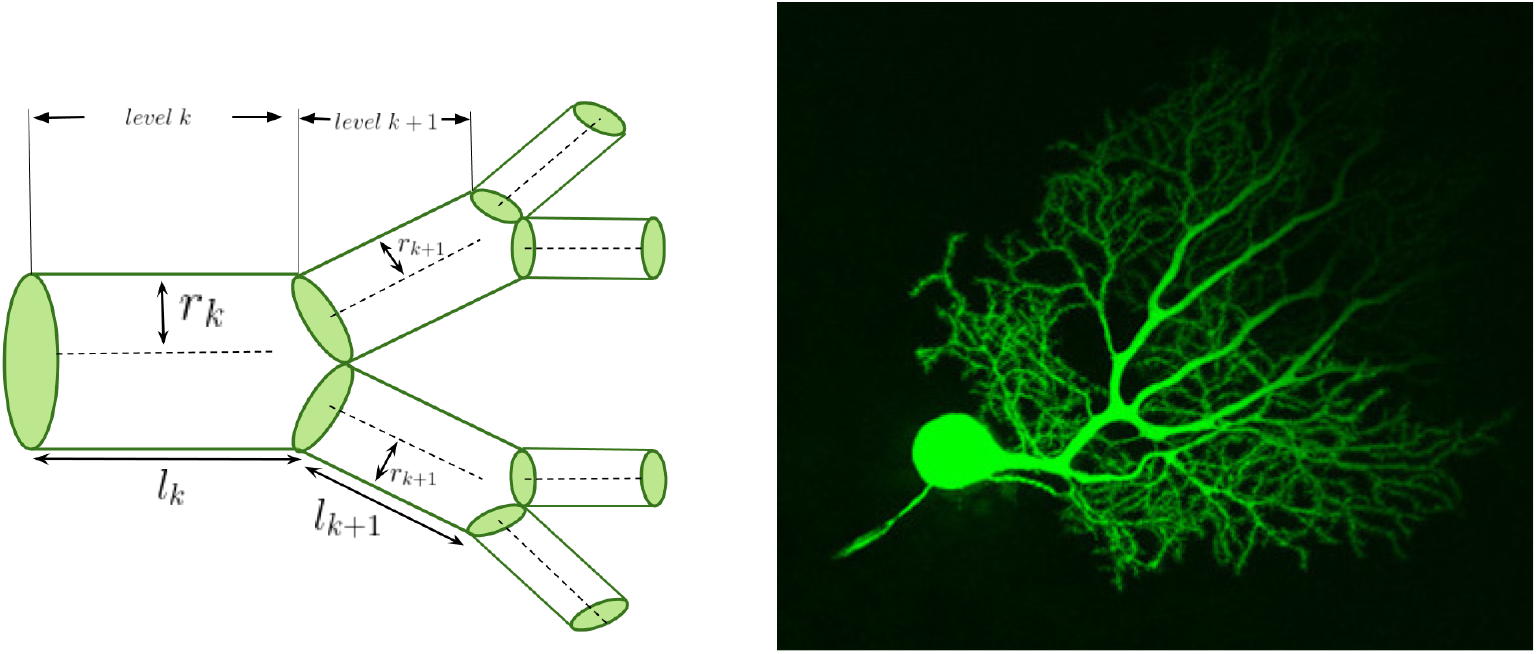
A hierarchical branching network. A visual depiction of the successive branching levels of a network and the quantities of interest alongside an image of a mouse cerebellar Purkinje neuron and its dendritic branching structure. This image was obtained using confocal microscopy and Lucifer yellow fluorescent dye. We have cropped this image available on CellImageLibrary.Org, distributed by Maryann Martone, Diana Price, and Andrea Thor [Martone et al., 2002].

In Equation 1, the first term is the power loss due to dissipation, given by 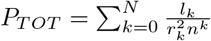. For a neuronal network, we define the power loss by the equation, 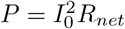, where *I*_0_ is the ionic current and *R_net_* is the resistance to current flow in the network. Axons and dendrites can be approximated as wires through which current flows and encounters resistance from the neuron fiber. The resistance is given by 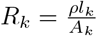, where *A_k_* is the cross sectional area of the wire, and *l_k_* is the length of the segment at that level. The parameter *ρ* is the intrinsic resistivity of the axon or dendrite, and we are assuming that *ρ* is constant, meaning that the material is uniform [Johnston and Wu, 1995]. Approximating axons and dendrites as cylinders, the cross sectional area is 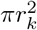 for level k, and the resistance is 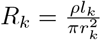. Following standard practice, we have absorbed all physical constants into the Lagrange constants, and the magnitude of these terms do not affect the theoretical predictions.

The second term represents conduction time delay, 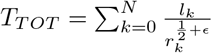, and arises because the average velocity of a signal along a single branch is 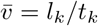, where *t_k_* is the time delay. We can solve this expression for *t_k_* and sum 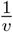 over the length of each branch [Ringo et al., 1994]. At each generation, we consider a single branch to denote the total path length of a signal, and we calculate the total conduction time delay by summing the time delays for single branches across all N generations. The parameter *ϵ* describes the degree of myelination. Previous work has shown that the conduction speed is proportional to the square root of the diameter for an unmyelinated fiber [Hodgkin, 1954], and proportional to the diameter for a myelinated fiber [Rushton, 1951]. Thus, an *ϵ* value of 0 corresponds to an unmyelinated fiber, and a value of 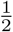 corresponds to a myelinated fiber.

We can switch between models that optimize either conduction time or power usage by varying *α* between 0 and 1, corresponding to the following two equations.

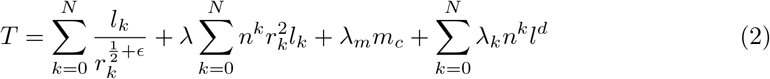

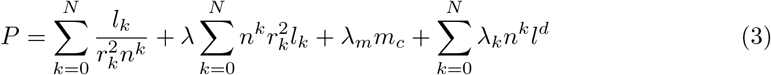

In these two equations, the governing optimization principle (first term) is constrained by brain region volume (second term), neuron size (third term), and space filling (fourth term). These quantities are held constant during the optimization. The last constraint comes from the fact that a resource distribution network must feed every cell in the body. Each branch in a given level of the network feeds a group of cells called the service volume, and the total service volume at each level is preserved. The service volumes vary in proportion to 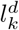, so the total volume is proportional to 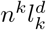 [Savage et al., 2008]. We assume that the branching ratio is constant, so the number of vessels at level k is *n^k^*. We can define the total volume as 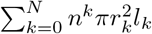, based on the assumption that the projections are cylindrical and branches are symmetric. We absorb the constant *π* into the Lagrange multiplier λ. The term that describes size, *m_c_*, is the mass of the cell. Although our formulation of energy consumption in this model relates to the power loss due to dissipation in signalling, a significant portion of energy consumption in neurons is involved in maintaining the resting membrane potential [Attwell and Laughlin, 2001]. This energy consumption is a per-volume quantity [Wang et. al., 2008], as it depends on the energy required by the sodium-potassium pump, which increases with increasing surface area of the neuron [Attwell and Laughlin, 2001]. Thus, the energetic cost of maintaining the resting membrane potential is captured in the total network volume.

In the section on the allometry calculation, we show that minimizing power (Eq. 3) subject to a conduction time delay constraint leads to a 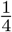-power scaling between conduction time delay and neuron size. This objective function can be described by the following equation:

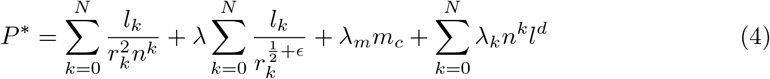

Equations 2, 3, and 4 are all specific cases of the more general Equation 1, with varying values of *α* as well as choice of constraint functions.

### Scaling Ratio Calculation

We use the method of Lagrange multipliers to solve for the values of the scaling ratios for radius and length, 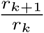 and 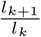 that minimize the objective function. Below, we show a sample calculation of the method of Lagrange multipliers for the case of power minimization (Eq. 3), where we choose the dimension d to be 3. A more detailed calculation can be found in Text S1.

We will first minimize P by differentiating with respect to radius at an arbitrary level *k*, setting the result equal to 0.

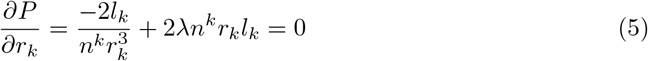

Solving for the Lagrange multiplier, we have:

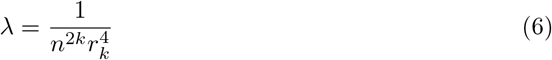

Using the fact that the Lagrange multiplier is a constant and thus the denominator must be constant across levels, we can solve for the scaling ratio:

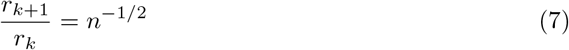

To find the length scaling ratio, we minimize P with respect to length at an arbitrary level *k*, and set the result equal to 0.

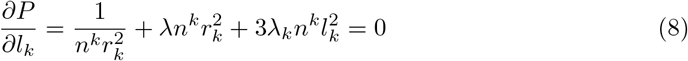

We solve for the Lagrange multiplier λ*_k_* by substituting λ, as calculated in (6). As before, using the fact that the denominator must be constant across levels and substituting in the scaling ratio in (7) for radius, we can solve for the scaling ratio for length:

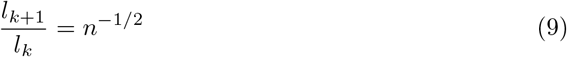

This method is used to solve for the scaling ratios for radius and length for the other cases and compared to empirical results. These findings are summarized in Table 1 in the Results section.

**Table 1.** Results for Radius Scaling Ratio Theoretical Predictions.

| Biophysical Principle | Prediction | Closest Biological Match | Data Mean |
| --- | --- | --- | --- |
| Time Minimization, Unmyelinated ($\mathbf{T}, \epsilon = \mathbf{0}$) | $n^{-2/5} \approx 0.76$ | Peripheral Nervous System Neurons | $0.76 \pm 0.008$ |
| Time Minimization, Myelinated ($\mathbf{T}, \epsilon = \frac{1}{2}$) | $n^{-1/3} \approx 0.79$ | Axons | $0.79 \pm 0.001$ |
| Power Minimization with Fixed Volume ($\mathbf{P}$) | $n^{-1/2} \approx 0.71$ | Purkinje cell dendrites | $0.69 \pm 0.007$ |
| Power Minimization with Fixed Time Delay ($\mathbf{P}^*$) | $n^{-2/3} \approx 0.63$ | Motoneuron dendrites | $0.63 \pm 0.007$ |

### Allometry Calculation

We now use the objective function P* (Eq. 4), to derive a functional scaling relationship between conduction time delay and body mass. Here, we consider the unmyelinated case (*ϵ* = 0), and the case of 3-dimensional space filling (d = 3).

We begin by setting the derivative of P* with respect to radius and length equal to zero to solve for the multipliers λ and λ*_k_*, respectively, at the stationary point. Substituting the expression for λ*_k_* back into the original expression for *P\**, we get an expression that simplifies to the original power term that it minimized, 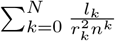. For simplicity, we replace the power term with *P* and the time delay constraint term with *T* and rearrange. This calculation is shown in detail in Text S2.

Previous results have shown a proportional relationship between *m_c_*, the mass of a single neuron, and the fourth root of an animal’s body mass, *M*^1*/*4^ [Savage et al, 2007]. Thus, we can replace this term and consider a new Lagrange multiplier with the absorbed constant:

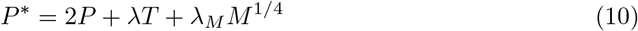

We will now take the derivative of this term with respect to M, the mass, and set it equal 0.

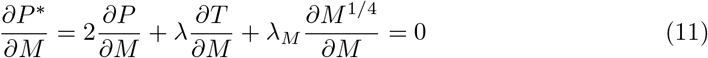

Previous results have shown that the energetic cost, which we have interpreted here as power loss due to dissipation, decreases with increasing body weight of animals at a linear rate [Wang et al., 2008]. Thus, we can express 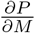 generally as a negative constant, −*C*. We can rewrite the above expression as follows:

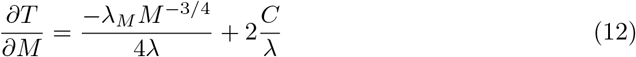

Solving and applying the initial condition that T=0 when M=0, we have:

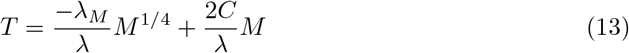

Thus, from this equation, we have extracted the scaling relationship, a mixed power law relationship including a 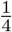-power law and a linear term with relative weights. Figure 5 shows experimental data that supports this theoretical result of the 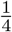-power law.

## Methods

To test the theoretical predictions and model, it is important to look at empirical data for scaling ratios for radius and length between child and parent branches in successive levels. We analyzed data from NeuroMorpho.Org - an online database with digital reconstructions from a wide range of species [Ascoli et al., 2007]. Figures 2,3, and 4 show five examples of images of neuron reconstructions obtained from NeuroMorpho.Org. These reconstructions are obtained by tracing neuron image stacks obtained using various microscopic and staining techniques for in vitro neurons and slicing at regular intervals. This database provides 3D reconstruction data that are organized in text files by pixels, in files that specify a pixel ID label for each point, the x,y,z spatial coordinates, the radius of the fiber at each point, and a parent pixel ID, referring to the adjacent pixel previously labelled. The scaling ratios for radius and length can be obtained by organizing this data in terms of branches. This is accomplished by finding the pixels at which the difference between the child pixel ID and the parent pixel ID is greater than 2, which can be defined as branching points. Based on the branching points, a branch ID and parent branch ID can be assigned to each of the pixels.

**Fig 2.**
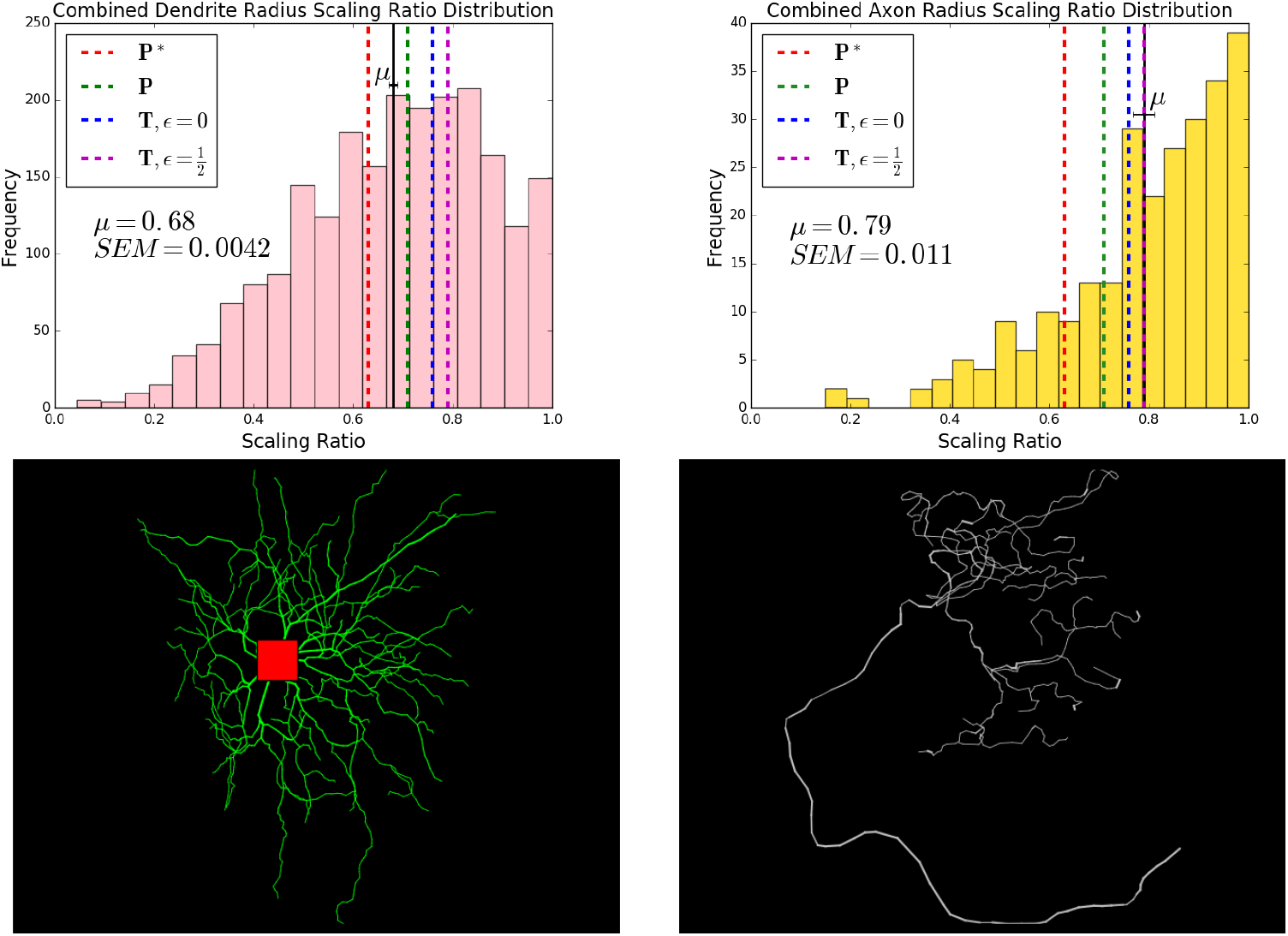
Comparison of Dendrite and Axon Radius Scaling Ratio Distributions, Combined. Histograms showing the distributions of radius scaling ratios for axons and dendrites combined from a range of species, brain regions, and cell types available on NeuroMorpho.Org. The mean dendrite scaling ratio is 0.68 ± 0.004 and the mean axon scaling ratio is 0.79 ± 0.001. In the figure, *μ* represents the mean and *SEM* represents the standard error of the mean (SEM). The standard deviations of the distributions are 0.19 for dendrites and 0.17 for axons. The black solid lines denote the mean in the distributions, shown with error bars, and the red, green, blue, and magenta dashed lines represent the theoretical predictions for various objective functions. The closest theoretical predictions for the dendrite scaling ratio mean are the optimal scaling ratios for function *P*, minimizing power with fixed volume, *n*−^1/2^ ≈ 0.71, and for function *P\**, minimizing power with fixed time delay, *n*−^2/3^ ≈ 0.63. The closest theoretical predictions for the axon scaling ratio mean are the optimal scaling ratios for function *T*, minimizing time delay, the myelinated case with 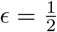, *n*−^1/3^ ≈ 0.79, and the unmyelinated case with *ϵ* = 0, *n*−^2/5^ ≈ 0.76. We restricted radius scaling ratio data to values that are less than 1.0. The representative reconstruction images show the characteristic differences in morphology between dendritic and axonal trees. The dendritic tree, shown on the left, is taken from an elephant cerebellar Golgi cell [Jacobs et al, 2014]. The axonal tree, with a representative long parent branch, is taken from a mouse touch receptor [Lesniak et al, 2014].

**Fig 3.**
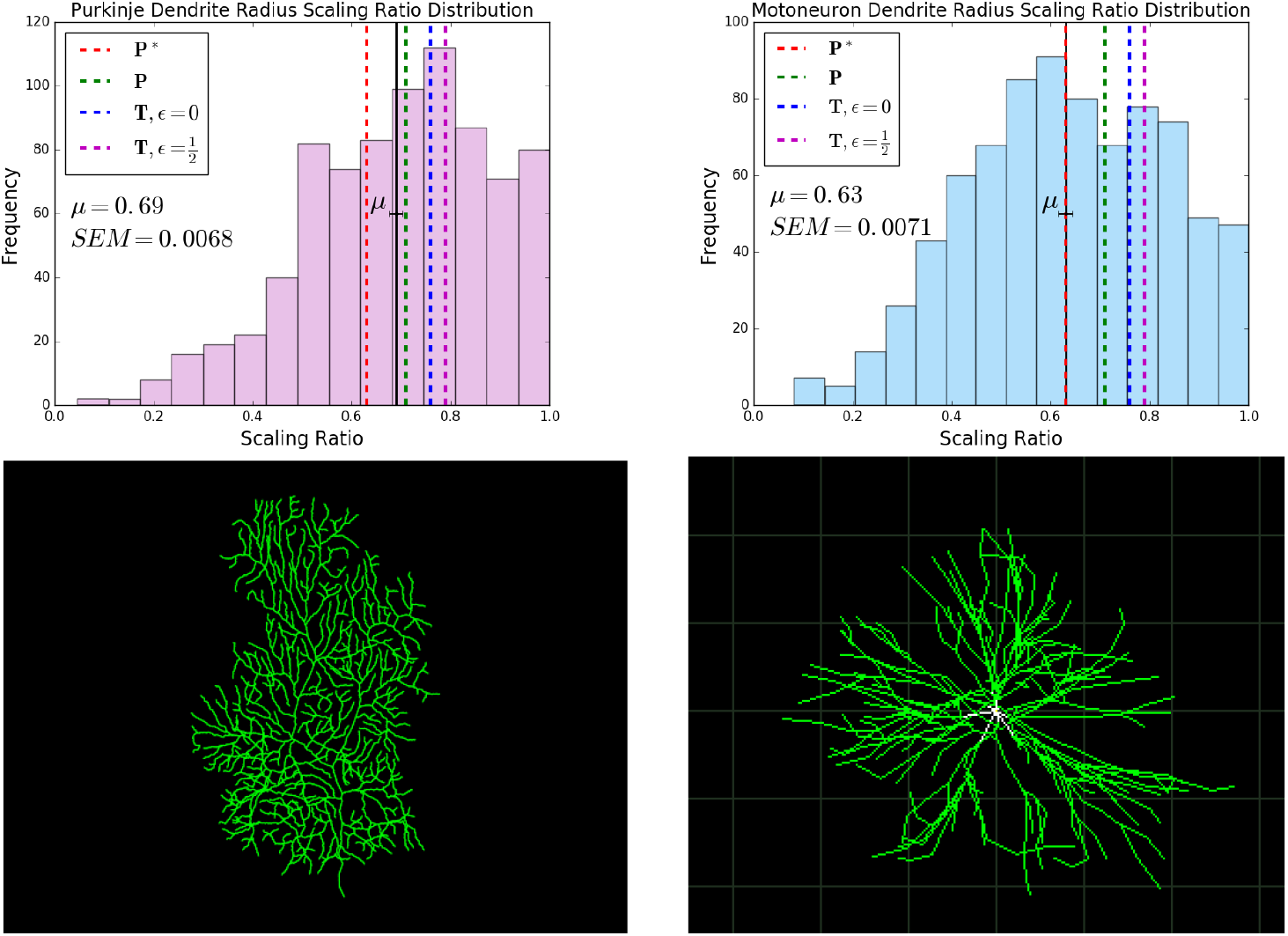
Comparison of Radius Scaling Ratio Distributions of Cerebellar Purkinje Cell and Motoneuron Dendrites. A comparison of histograms showing the distribution of radius scaling ratios observed in dendrites of Purkinje cells and motoneurons, along with representative images. For Purkinje cells, we observe an average radius scaling ratio of 0.69 ± 0.01, and for motoneurons, we observe an average radius scaling ratio of 0.63 ± 0.01. In the figure, *μ* represents the mean and *SEM* represents the standard error of the mean. The standard deviations of the distributions are 0.19 for Purkinje Cells and 0.20 for motoneurons. We have restricted radius scaling ratio data to values that are less than 1.0. The black solid lines denote the mean values in the distributions, shown with error bars, and the red, green, blue, and magenta dashed lines represent the theoretical predictions for various objective functions. The closest theoretical prediction for Purkinje cells is the optimal scaling ratio for function *P*, minimizing power with fixed volume, *n*−^1/2^ ≈ 0.71. The closest theoretical prediction for motoneurons is the optimal scaling ratio for function *P\**, minimizing power with fixed time delay, *n*−^2/3^ ≈ 0.63. The representative image for the Purkinje cell is from a mouse [Murru et al., 2019] and the representative image for the motoneuron is from a cat spinal motoneuron [Cullheim et al., 1987].

**Fig 4.**
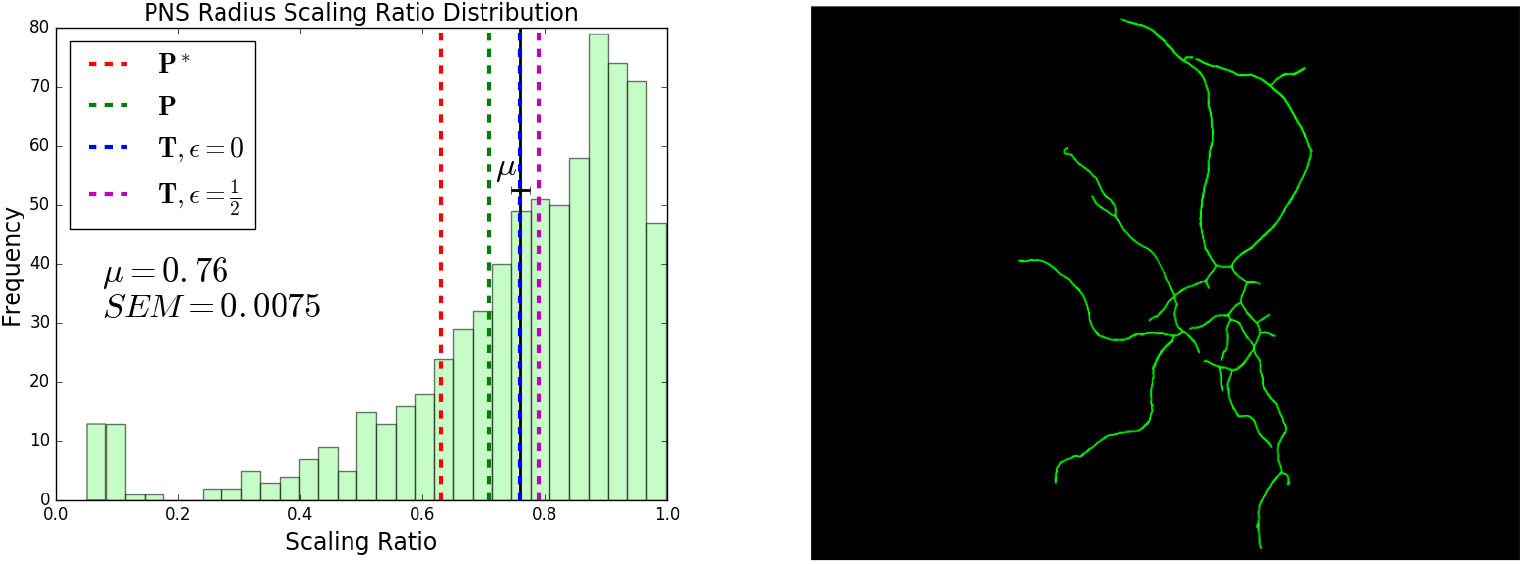
Peripheral Nervous System Neurons. A histogram showing the distribution of radius scaling ratios in Peripheral Nervous System (PNS) neurons, along with a representative image of the dendritic tree of a mouse sensory neuron [Shevalye et al., 2015]. We observe an average radius scaling ratio of 0.76 ± 0.01. In the figure, *μ* represents the mean and *SEM* represents the standard error of the mean. The standard deviation of the distribution is 0.20. We have restricted radius scaling ratio data to values that are less than 1.0. The black solid lines denote the mean in the distributions, shown with error bars, and the red, green, blue, and magenta dashed lines represent the theoretical predictions for various objective functions. The closest theoretical prediction is *n*−^2/5^ ≈ 0.76, the optimal scaling ratio for the function *T* that minimizes time delay for unmyelinated fibers, *ϵ* = 0.

The radius can be extracted from each of the branches by taking each of the radius values in each branch and averaging them by the following formula, defining each branch as branch k, where the pixels i range from 1 to *N_k_*, where *N_k_* is the last pixel of each branch:

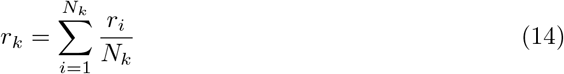

The length of each branch can be extracted by summing up the Euclidean distances between each of the points in the branch by the following formula:

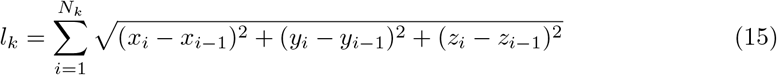

Once the radius and length of each of the branches is found, the scaling ratios are computed by dividing the daughter radius or length, respectively, by the corresponding value for the parent branch. Through this method and using the Python library matplotlib, we generate histograms to visualize the distributions. For the radius distributions, we find a large peak at 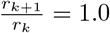, which is likely due to the resolution limit of the images. After a certain level, the radius for each of the branches is equivalent to the pixel size itself. Thus, in our distributions for radius, we focused on the data for scaling ratios that are less than 1.0. We use solid black lines to denote the mean values in the data, and error bars represent twice the Standard Error of the Mean (SEM), the standard deviation divided by the square root of the number of data points.

We look at neuron reconstructions from both axons and dendrites, and from a range of cell types, brain regions, and species. More detailed information about the source of each of the individual reconstructions can be found in Text S4.

For dendrites, we looked at three different types of cells: Golgi cells, Purkinje cells, and motoneurons. The Golgi cells are from *Giraffa*, *Homo Sapiens*, *Loxodonta africana*, *Megaptera novaeangliae*, *Neofelis nebulosa*, *Pan troglodytes*, *Panthera tigris* [Jacobs et al., 2014], and *Mus musculus* [Vervaeke et al., 2012]. The Purkinje cells are from *Cavia porcellus* [Rapp et al, 2994], *Mus musculus* [De Munter et al., 2016, Chen et al., 2013, Murru et al., 2019, Martone et al., 2003], and *Rattus* [de Luca et al., 2009, Martone et al., 2003, Vetter et al., 2001]. The motoneurons are from *Danio rerio* [Morrice et al., 2018, Svahn et al., 2018], *Felis Catus* [Cullheim et al., 1987], *Mus musculus* [Leroy et al., 2014], *Oryctolagus cuniculus* [Steele et al., 2020], *Rattus* [Rotterman et al., 2014], and *Testudines* [Chmykhova et al., 2008]. In Figure 2, we look at the combined dendrite data for all cell types and species. In Figure 3, we look at the radius scaling ratios of Purkinje cells and motoneurons individually, and draw comparisons between the two.

Due to the small size of axons and the limited resolution of images, the data available on NeuroMorpho.Org are limited in scope. The data shown in Figure 2 was taken from the following species: *Anisoptera* [Gonzalez-Bellido et al., 2013], *Brachyura* [Bengochea et al., 2018], *Drosophila melanogaster* [Jefferis et al., 2007], *Gallus gallus domesticus* [Garrido-Charad et al., 2018], and *Rattus* [Martins et al., 2017]. The neurons were taken from a range of brain regions: the midbrain, the hippocampus, the antennal lobe, the optic lobe, and the ventral nerve cord.

To study peripheral nervous system neurons, we sampled from reconstruction data that was labelled by region on NeuroMorpho.Org. This data, shown in Figure 4, was taken from *Drosophila melanogaster* [Herman et al., 2018, Nanda et al., 2018, Ye et al., 2011] and *Mus musculus* [Badea et al., 2012, Lesniak et al., 2014, Canavesi et al., 2020, Shevalye et al., 2015] and includes dendritic arborizations, sensory neurons, somatic neurons, and touch receptors.

To look at functional scaling relationships between mass and conduction time delay, we first look at data for conduction time delay in motoneurons and sensory neurons across a range of species sizes, listed in order of size: *Soricidae*, *Mus musculus*, *Rattus*, *Cavia porcellus*, *Oryctolagus cuniculus*, *Felis Catus*, *Canis lupus familiaris*, *Sus scrofa*, *Ovis aries*, *Giraffa*, and *Loxodonta africana* [More and Donlean, 2018]. Using the mean conduction velocity measured in studies of each species, this conduction time delay data was calculated by considering the animal leg length predicted from the average body mass. We use a log-log plot, shown in Figure 5, to obtain a power law relationship between body mass and conduction time, where the slope is equal to the power.

**Fig 5.**
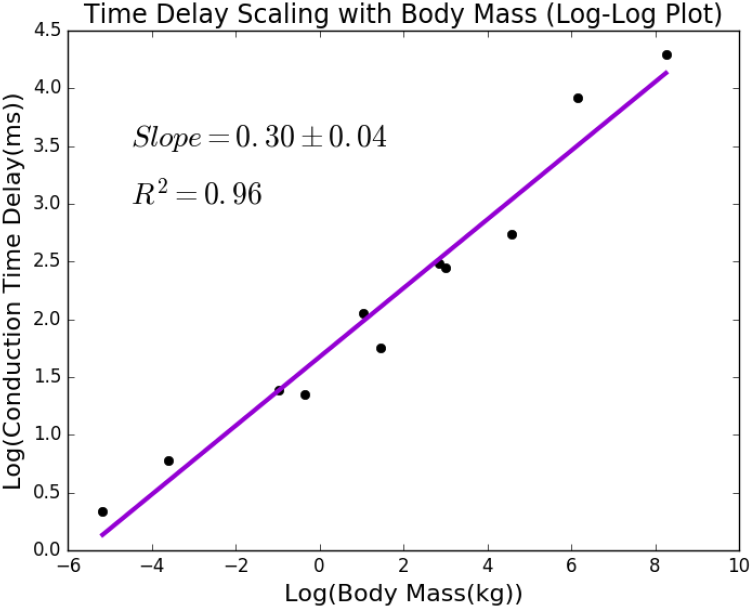
Scaling of Conduction Time Delay and Species Mass. A scatter plot showing the relationship between the log of the conduction time delay and the log of the body mass of a range of species. Here, the slope, 0.30 ± 0.04, corresponds to the power that relates species mass to conduction time delay. This is close to our theoretical result of 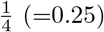.

## Results

We compared theoretical predictions for scaling ratios calculated from objective functions *T*, *P*, and *P\** with the mean values we measure from the data. As mean values capture the average overall branching properties for axons and dendrites, the mean represents the most natural and straightforward staring point for comparing our general theory with empirical data. As theory is refined and additional predictions are made, other features of the distribution should also be measured and compared (see Discussion section). Based on the results of these comparisons for different types of neurons and processes, we determined the functional properties that play the greatest role in determining structure for different processes and cell types.

### Theoretical Predictions

Using the model and the method of undetermined Lagrange multipliers as detailed above, we made theoretical predictions for functions using different values of the parameters. Table 1 shows the results for the various objective functions minimizing conduction time delay and power. The approximations listed are based on the simplifying assumption that the network is purely bifurcating, with a branching ratio of 2.

We consider the theoretical predictions for four objective functions. The first two objective functions are specific cases of *T* (Eq. 2), minimizing conduction time delay. We consider this function for two possible values of the parameter *ϵ*. The unmyelinated case corresponds to *ϵ* = 0, and 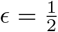 signifies the myelinated case. The second two objective functions are minimizing power. The objective function *P* (Eq. 3) minimizes power subject to volume, mass, and space filling constraints, where volume is fixed. The alternative objective function *P\** (Eq. 4), is the objective function minimizing energy subject to conduction time, mass, and space filling constraints, where time delay is fixed. For power minimization, we focused on the unmyelinated case where *ϵ* = 0, since its predictions align most with the data from dendrites, which are typically unmyelinated.

For all of the calculations, we considered different values of the parameter d, the dimension of space filled by the processes. A value of d=2 signifies neuron processes that branch into a 2-dimensional plane, such as Purkinje cells in the cerebellum. A value of d=3 signifies neuron processes that fill a 3-dimensional space, such as motoneurons [Squire et al., 2013]. We interpreted the volume constraint as a material constraint, assuming that the processes are cylindrical for both 2- and 3-dimensional space-filling. It is interesting to note that the dimension of space filling does not affect results for radius scaling ratios. However, it does play a role in the results of the theoretical predictions for length, as we have detailed in Table S1 and Table S2. Given the lack of agreement between the theoretical predictions for length scaling ratios and the data, we focused on radius scaling ratios in this analysis.

### Dendrites and Axons

Figure 2 shows histograms that illustrate the differences in distributions of radius scaling ratios for dendrites and axons, along with representative images of the morphology of these two processes. Axons generally carry signals from the cell body to the synapses, where they transfer information to the dendrites of other neurons. Dendrites have extensive, tree-like structures and generally connect with the axons of other neurons to carry signals to the cell body. The distributions observed for these scaling ratios resemble the distributions observed in scaling ratios of cardiovascular networks, with the radius scaling ratios exhibiting a normal distribution.

In this figure, we show the comparison of the mean dendrite radius scaling ratio, 0.68 ± 0.004, with theoretical predictions from the four different calculations. We find that the dendrite radius scaling ratio mean is closest to the theoretical predictions from the objective functions minimizing power. The mean lies in between the optimal scaling ratios for function *P*, so *n*−^1/2^ ≈ 0.71, which holds volume to be fixed, and function *P\**, so *n*−^2/3^ ≈ 0.63, which holds time delay to be fixed. Later, in Figure 3, we looked at the distributions of radius scaling ratios in these Purkinje cells and motoneurons individually and compared them to the closest theoretical results individually.

Note that the radius scaling ratio mean for axons, 0.79 ± 0.001, is significantly larger than the mean radius scaling ratio observed for dendrites, 0.68 ± 0.004. The axon scaling ratio mean in the data is closest to the theoretical prediction, *n*−^1/3^ ≈ 0.79, for the objective function that minimizes time, *T*, for myelinated fibers, 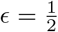. The next closest prediction, *n*−2/5 ≈ 0.76, is that of the objective function that minimizes time, *T* for unmyelinated fibers, *ϵ* = 0. This suggests that time minimization and myelination are important factors that determine the structure for axons.

### Purkinje Cells and Motoneurons

One of the parameters that we built into our theoretical model is *d*, the dimension of space filling of the processes. Thus, we looked at the comparison of results from data from representative cells with 2-dimensional and 3-dimensional dendritic trees. For dimensional dendritic trees, we looked at cerebellar Purkinje cell data from rodents including mice, rats, and guinea pigs. For 3-dimensional dendritic trees, we looked at motoneurons from a range of species including rodents, amphibians, cats, and humans. The histograms for these two cell types, along with a representative image for each type, are shown in Figure 3. The theoretical results of minimizing power and time cost functions while varying the parameter *d* do not capture the differences in radius scaling ratios observed in this data. We hypothesized that the differences observed can be explained by other principles such as the functional differences of these cell types. The mean for Purkinje cells, 0.69 ± 0.01, agrees with the theoretical predictions for the function *P*, power minimization with a volume constraint, *n*−^1/2^ ≈ 0.71, while the mean for motoneurons, 0.63 ± 0.01, agrees with the theoretical predictions for function *P\**, power minimization with a time constraint, *n*−^2/3^ ≈ 0.63. Based on the results of the comparison of Purkinje cells and motoneurons, we concluded that volume plays a greater role in constraining the structural design of Purkinje cells, while time plays a greater role in constraining the structural design of motoneurons.

### Peripheral Nervous System Neurons

In the Peripheral Nervous System (PNS), motoneurons play an important role in the exchange of information with sensory neurons. Peripheral nerves carry sensory information and interact with motoneurons, which directly innervate effector cells such as muscles [Squire et al., 2013]. Thus, the importance of conduction time as a constraint for motoneurons motivated us to examine data from other types of PNS neurons such as sensory neurons. Figure 4 shows the radius scaling distribution of a sample of the PNS neurons labelled by region on NeuroMorpho.Org. This data was taken from flies and mice. The mean radius scaling ratio, 0.76 ± 0.01, is closest to the theoretical prediction, *n*−^2/5^ ≈ 0.76, for the objective function, *T*, that minimizes time for unmyelinated fibers, *ϵ* = 0. This suggests that time is an important factor in optimizing structure for PNS neurons.

### Time Delay Scaling

So far, we have focused on predictions and data combined from species of a range of sizes. Here, we consider how function varies across species of a range of body masses. We used *P\**, the equation minimizing power with fixed time delay. As shown in the Theory section, our theoretical calculations have led to the following relationship between conduction time delay and mass:

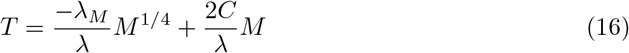

In order to test this theoretical result, we analyzed experimental data to determine an observed relationship between time delay as a function of species size. Previous experimental studies have looked at conduction time delay across species ranging from shrews to elephants [More and Donlean, 2018]. A regression analysis of the data shows that the 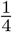-power mass term is more significant than the linear term, as is shown in more detail in Text S3. Furthermore, we used a log-log plot to determine the power of the relationship, plotting the log of the conduction time delay data against the log of the average body mass of each species. This plot is shown in Figure 5.

Our theoretical predictions suggest the presence of a 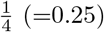 power law that relates species mass to neuron conduction time delay. These experimental results support this power law, as the power law determined from the data is 0.30 ± 0.04. The available data focuses on a limited range of masses, and it is possible that at a wider range of masses, a scaling law closer to the linear relationship might be observed. Further data and analysis of the relationship between the size of individual neurons and processes and species mass and between conduction velocity and time delay will provide useful insight into this allometry.

## Discussion

A comparative analysis of the radius scaling ratios of different processes and cell types suggests that there are selection pressures for different functional roles that underlie the diversity in neuron branching patterns. There are a number of characteristic differences observed between axons and dendrites. Axons are long and function to transmit signals over large distances, sometimes between different regions of the nervous system [Rall, 1977]. Moreover, axons have the unique property of myelination, which provides an important role in information transfer in the nervous system [Waxman and Swadlow, 1977]. Our results indicate that the radius scaling ratio mean for axons is closest to the prediction that minimizes time for conduction through myelinated fibers, which supports this notion that information processing speed is a key principle governing the structure of axons.

In contrast, dendritic trees are relatively short, have more extensive branching, and generally do not conduct action potentials [Rall, 1977]. Previous theoretical work on wiring optimization in cortical circuits similarly proposes that there are differing evolutionary selection pressures governing axons and dendrites. Rather than conduction time delay, the key principle behind dendritic structure is passive cable attenuation [Chklovskii, 2000]. Our results suggest that dendrites are optimized to minimize power, which is related to a voltage drop, with a volume constraint that we have interpreted as a cost in materials. Thus, minimizing power in our theoretical framework is effectively minimizing the attentuation of the passive signals in dendrites.

There is a great deal of diversity in the branching structures of dendritic trees, and the differences in scaling ratio distributions among the different types gives us important insights into their distinct functional roles. We found that the structure of Purkinje cells and motoneurons are both governed by power minimization, and Purkinje cell structure is constrained by volume while motoneuron structure is constrained by time delay. The predictions and results from the data for Purkinje cells and motoneurons are supported by previous theoretical and experimental results [Hillman, 1979]. We conclude that time plays a greater role in optimizing the structure for motoneuron dendrites.

Efficiency in information processing is a key function of neurons in the sensorimotor system, and our results emphasize that function as a key feature governing their structural design. When organisms are exposed to environmental stimuli, it triggers a response in the motor system that must be executed very rapidly. Some of these responses are innate, and some are learned through practice, gradually increasing in speed [Vidal et al., 2015]. We found that structure of neurons in the peripheral nervous, such as the sensory neurons that relay information from the environment to motoneurons, is governed by time minimization, which is consistent with the evolutionary function of the sensorimotor system. The correspondence of our theoretical predictions with empirical measurements from neurons of different types supports intuitive notions about neuron computation in these specific cell types.

Major steps have been taken to advance and to quantitatively formalize the laws first set forth by Santiago Ramon y Cajal for how functional principles dictate neuron morphology. Notably, Hermann Cuntz’s group quantifies Ramon y Cajal’s laws of conservation of time and material using principles from graph theory to computationally generate biologically accurate axonal and dendritic trees [Cuntz et al., 2010]. Furthermore, Dmitri Chklovskii formalizes the differences in structure and function between axons and dendrites and even considers the role of cable diameter in wiring optimization [Chklovskii, 2004]. Also of note is that material constraints related to increasing diameter play a role in limiting the scaling of conduction speed in larger animals, leading to longer delays [Ringo et al., 1994]. Finally, more recent work by Samuel Wang, Simon Laughlin, and Terrence Sejnowski considers how energy consumption constrains the design of neuronal networks, particularly when considering differences across species of different sizes [Laughlin and Sejnowski, 2003, Wang et al., 2008]. Here, we have integrated results from these studies and provided a volumetric explanation of these branching structures in terms of their biological and physical function across scales that considers conduction speed, material costs, metabolic costs, and space-filling. The correspondence between theoretical predictions and empirical measurements of radius scaling ratios in neuron branching processes provides important insights into the relationship between structure and function.

So far, we have looked at optimization problems minimizing power and time individually. However, it is possible that there might be intermediate values, and different cell types might have different relative importance of time and power in determining structure. A possible avenue for future work is using numerical methods to extend the number of functional principles we consider in each prediction. This might provide a more biologically useful estimate for scaling ratios, as it is likely that neuron cell structures are designed to optimize not only conduction speed or energy efficiency, but a relative combination of both.

The similarities in distributions of scaling ratios in radius and length between neurons and cardiovascular networks suggest that a unifying framework underlies these diverse biological systems. Although the WBE framework for cardiovascular networks provides us with a solid framework to build upon in analyzing neuronal networks, there is still much work to be done in adapting this model for neurons. It is widely understood that the morphology of dendritic arbors are not static, but are constantly modifying based on interactions with surrounding neurons and glia [Squire et al., 2013]. Incorporating this dynamical aspect of neuron morphology will be useful in future development of our model. Moreover, we have formulated the space filling constraint based on the idea that cardiovascular networks are optimized such that vessels feed every cell in the body. However, neurons exhibit more complex space filling patterns new to their interactions with one another, such as tiling and self-avoidance [Cameron and Rao, 2010]. It might also be fruitful to consider different formulations of the space filling constraints for different types of neurons. For example, axons tend to have a projections featuring a longer parent branch, and the daughter branches occur further away from the soma. Indeed, previous work has extended the WBE model to look at scaling in plants [Price and Enquist, 2007]. We might look into applying previous work on space filling for plants such as palm trees, which have similar morphology.

Additionally, we have represented the energy consumption here as the power lost due to dissipation during signalling. In neurons, however, maintaining the resting membrane potential makes up a significant fraction of the energetic costs. Here, we assumed that this cost is captured in the volume term in the model. However, it might be possible to more explicitly formalize the inclusion of the resting potential via the incorporation of additional factors that affect this cost. For example, myelination effects the capacitance of axons, and the energy required to maintain the resting potential varies linearly with capacitance [Wang et al., 2008]. Incorporating these complexities in our model might improve its biological accuracy and usefulness when comparing predictions to empirical data from neurons.

Another future direction is to employ alternate labeling schemes for the branching levels in order to extract more meaningful results for length scaling ratios, we might look into applying alternate labeling schemes for the branching levels. Previous work on river networks has used an alternative labeling scheme - called Horton-Strahler labeling - where the first level begins at the tips, and higher levels are determined when two branches of the same level combine. This has been applied to other networks in biology [Turcotte et al., 1998]. Hermann Cuntz’s group has also applied this ordering method to analyze dendritic trees, finding differences in branching metrics across neuron cell types [Vormberg et al., 2017]. We hypothesize that applying this labeling scheme to define branching levels for length will give a distribution of scaling radios that looks more like the normal distributions observed for radius scaling ratios, and means that agree more closely with our theoretical predictions. This is a major goal of our future work, both for neurons and cardiovascular networks. Moreover, our analysis of the comparisons of the theoretical predictions to the data involves simply information about the mean values. We chose to look at the mean rather than the mode of the distributions to take into account the spread of the distributions, but these values do not align in all cases. Future work might look further into additional features of the distributions of radius and length scaling ratios in order to extract more information from the data.

Throughout this model, we have assumed that branching is symmetric - the radius and length of daughter branches are identical. Previous work has attempted to capture asymmetry in cardiovascular networks and plants [Brummer et al., 2017]. Another major goal of our future work is applying this theoretical framework to look at branching of neuron processes, and using branching properties related to asymmetry to compare different cell types.

Beyond the scaling ratios for successive branches in the individual neuron processes, it is interesting to consider allometric scaling relationships of species size and functional properties. Previous work on cardiovascular networks has extracted an allometric scaling relationship that relates species size (or mass) with volume [Savage et al., 2008]. Moreover, previous work on scaling has shown an allometric scaling relationship between single cell neurons and animal body mass [Savage et al, 2007], and when brains grow in size, they require more extensive axonal trees to traverse greater distances [Bekkers and Stevens, 1990]. Building on these ideas from our theoretical formulation of the objective function that minimizes power subject to the constraint of fixed conduction time delay, we were able to extract a functional scaling relationship between species size and time delay for unmyelinated fibers. We derived that there is a mixed power law relationship between animal body mass and conduction time delay, including both a term with a 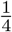-power and a linear term. The presence of the 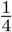-power law is supported by experimental data of conduction time delay from species of a range of masses: the conduction time delay scales with the fourth root of the animal body mass.

An interesting aspect of this result is that neurons in larger animals have longer conduction delays. These results are important to consider in the context of evolution - longer delays might provide a functional explanation for the increased specialization of brain function hemispheres. Due to the greater conduction time delays, it might be advantageous for larger brains to exhibit more specialization, and organize cells with information about related memories and skills in localized clusters [Ringo et al., 1994], thus improving the efficiency of information processing.

We conclude that neuron function places profound constraints on neuron morphology, thus cementing the foundations in Ramón y Cajal’s documentation and resulting theoretical and computational formalism proposed by Cuntz and Chklovskii, and extending it to include metabolic constraints and consider the volumetric aspect of morphology. Our modern approach provides a technologically sophisticated way to measure and quantify neuron morphology, and a mathematically and theoretically advanced way to describe the influence of biophysical constraints in selecting morphological patterns in neurons. Combining empirical measures with our theoretical predictions, we showed fundamental differences between axons and dendrites and between Purkinje cells and motoneurons in ways that in turn depend on the myelination of axons and the dimension of space being filled by the branching processes. Future work in this direction will shed even more light on these foundational questions by obtaining larger amounts of data at higher resolutions across more species and more cell types. Indeed, looking across species and cell types will also help reveal further differences in neuronal function and tradeoff among different principles that may transform how we understand the function and form of the brain.

## Supporting Information

**Figure S1** Length Scaling Ratio Distributions for Dendrites and Axons

**Table S1** Length Scaling Ratio Theoretical Predictions from Time Minimization

**Table S2** Length Scaling Ratio Theoretical Predictions from Power Minimization

**Text S1** Scaling Ratio Calculation

**Text S2** Allometry Calculation

**Text S3** Allometric Scaling Relationship Regression Analysis

**Text S4** Overview of Reconstruction Data Sources

## Acknowledgments

This material is based upon work supported by the National Science Foundation Graduate Research Fellowship under Grant No. (NSF grant number DGE-1650604 and DGE-2034835) and the National Institutes of Health Systems and Integrative Biology Training Grant under Grant No. (NIH grant number 2T32GM008185-31 and 5T32GM008185-32). Any opinions, findings, and conclusions or recommendations expressed in this material are those of the authors and do not necessarily reflect the views of the National Science Foundation or the National Institutes of Health.

